# Recursive splicing is a rare event in the mouse brain

**DOI:** 10.1101/2020.09.10.291914

**Authors:** Sohyun Moon, Ying-Tao Zhao

## Abstract

Recursive splicing (RS) is a splicing mechanism to remove long introns from messenger RNA precursors of long genes. Compared to the hundreds of RS events identified in humans and drosophila, only ten RS events have been reported in mice. To further investigate RS in mice, we analyzed RS in the mouse brain, a tissue that is enriched in the expression of long genes. We found that nuclear total RNA sequencing is an efficient approach to investigate RS events. We analyzed 1.15 billion uniquely mapped reads from the nuclear total RNA sequencing data in the mouse cerebral cortex. Unexpectedly, we only identified 20 RS sites, suggesting that RS is a rare event in the mouse brain. We also identified that RS is constitutive between excitatory and inhibitory neurons and between sexes in the mouse cerebral cortex. In addition, we found that the primary sequence context is associated with RS splicing intermediates and distinguishes RS AGGT site from non-RS AGGT sites, indicating the importance of the primary sequence context in RS sites. Moreover, we discovered that cryptic exons may use an RS-like mechanism for splicing. Overall, our findings provide novel insights into RS in long genes.

## Introduction

Recursive splicing (RS) is a splicing mechanism that removes a long intron into several smaller segments as opposed to in a large single unit(1–11). RS is untraceable in the mature messenger RNA (mRNA), and the direct evidence of RS is the splicing intermediates. However, RS splicing intermediates are unstable, making them difficult to be captured and analyzed. Whole-cell ribosomal RNA-depleted total RNA sequencing (total RNA-seq) and nascent RNA-seq have been used to identify RS splicing intermediates(3, 5, 7–9, 11).

RS has been widely studied in humans. Two earlier studies identified five and nine RS events in humans(3, 5), suggesting that RS might be a rare splicing mechanism for a small group of long introns. Intriguingly, a recent study using nascent RNA-seq identified 342 candidate RS sites in three human cell lines(9). Similarly, another study using nascent RNA-seq reported 5,468 RS events in a human cell line(11). These two studies suggested that RS might be a widely used splicing mechanism in mammals. However, given these extensive studies of RS in humans, only ten RS events have been reported in mice so far. Thus, the extent to which RS is used for intron splicing in mice remains largely unexplored.

Here, we report that nuclear total RNA-seq is enriched for RS splicing intermediates and nascent transcripts, which suggests that it is an efficient approach to investigate RS events. Using nuclear total RNA-seq data generated from the mouse brain, we identified novel RS events, examined the cell-type and sex specificity of RS, and analyzed RS splicing intermediates. We found that RS is a rare process of intron splicing in the mouse brain. We also found that some cryptic exons use an RS-like mechanism for splicing. Together, our findings provide novel insights into the mechanisms of RS in long genes.

## Results

### Nuclear total RNA is enriched for nascent transcripts and RS splicing intermediates in the mouse brain

Compared to hundreds of RS events identified in humans, only nine RS events have been reported in mice(5). Because RS tends to occur in long genes, to further investigate RS in mice, we analyzed RS in the mouse brain, a tissue that is enriched in the expression of long genes. To determine whether nascent transcripts and RS splicing intermediates are enriched in nuclear RNA (Fig. 1a), we analyzed nuclear ribosomal RNA-depleted total RNA sequencing (total RNA-seq) and whole-cell total RNA-seq data that we recently generated from the mouse cerebral cortex(12, 13) (Supplementary Figure S1a). As a control data set, we also analyzed poly(A) enriched messenger RNA-seq (mRNA-seq) data that were generated from the same mouse brain region(14).

**Fig. 1.**
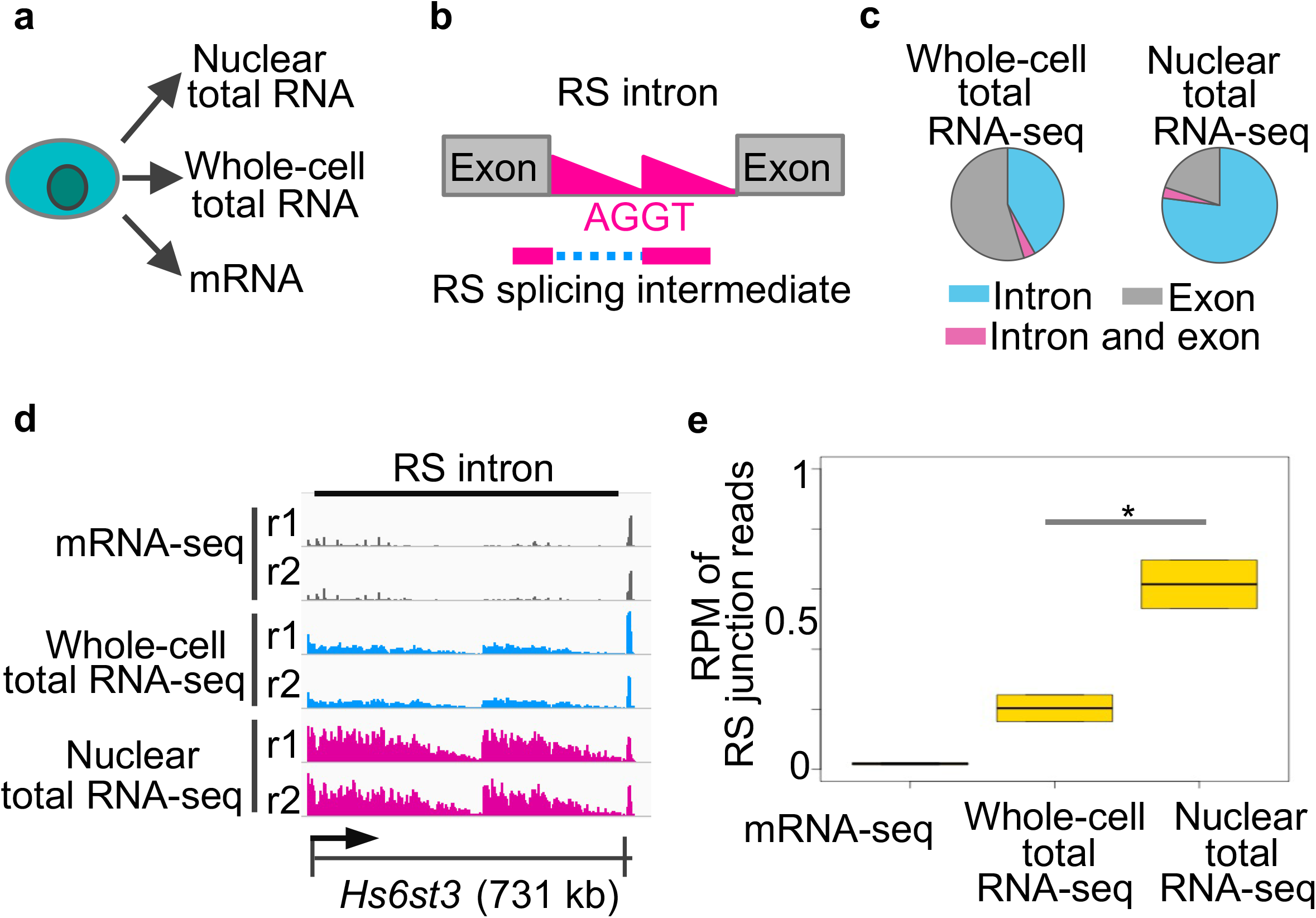
Nuclear total RNA is enriched for nascent transcripts and RS splicing intermediates in the mouse brain. **(a)** Schematic of the isolation of different types of RNA. **(b)** Schematic of the two features of RS, the saw-tooth pattern (red triangles) and the RS splicing intermediate. **(c)** Pie charts of loci of uniquely mapped reads in gene regions. **(d)** Sequencing profile at *Hs6st3*. r1, replicate 1. kb, kilobases. **(e)** Boxplot of normalized numbers of RS junction reads at *Hs6st3* RS site. RPM, reads per million uniquely mapped reads. *, *P* = 0.03, one-tailed t-test.

Nascent transcripts, which can be indicated by the sequencing reads mapped to introns(15), are important to identify RS events, because RS introns exhibit a saw-tooth pattern of nascent transcript signals from total RNA-seq data(3, 5) (red triangles, Fig. 1b). To determine the enrichment of nascent transcripts in nuclear RNA, we calculated the proportions of reads mapped to introns in nuclear total RNA-seq data, whole-cell total RNA-seq data, and mRNA-seq data. We found that 77% of the uniquely mapped reads from the nuclear total RNA-seq data are localized in introns, which is significantly higher than the 41% from the whole-cell total RNA-seq data and the 23% from the mRNA-seq data (*P* < 2.2^-16^, one-tailed Fisher’s Exact Test) (Fig. 1c and Supplementary Figure S1a). In addition, at a known RS intron in *Hs6st3*, we observed a more distinct saw-tooth pattern in the nuclear total RNA-seq data than in the whole-cell total RNA-seq data (Fig. 1d). These results suggest that nuclear total RNA is enriched for nascent transcripts.

RS splicing intermediates, which can be indicated by the junction reads spanning the upstream exon and the RS AGGT site (RS junction reads, Fig. 1b), is also important to identify RS events. To examine RS splicing intermediates in nuclear RNA, we calculated the numbers of junction reads at a known RS site in *Hs6st3*. We found that the number of RS junction reads at *Hs6st3* is 3-fold higher in nuclear total RNA-seq data than in whole-cell total RNA-seq data (Fig. 1e). Together, these results indicate that nuclear total RNA is enriched for nascent transcripts and RS splicing intermediates.

### Identify RS sites using nuclear total RNA-seq data

To identify RS sites from nuclear total RNA-seq data, we modified a previously developed pipeline and incorporated additional criteria(5) (Fig. 2a). First, we extracted all junction reads spanning the AGGT sites in the mouse genome and then focused on the AGGT sites located in introns that are longer than 1 kb. To exclude unannotated exons from our analyses, we selected AGGT sites that show more than two-fold enrichment of RS junction reads in nuclear total RNA-seq than in mRNA-seq. To increase the degree of confidence, we selected sites containing 10 or more RS junction reads as RS site candidates. To enhance the detection of RS junction reads, one of the nuclear total RNA-seq sample was sequenced to extra depth, resulting in 428 million uniquely mapped reads (Supplementary Figure S1a). Lastly, we visually inspected the candidates to select sites that show saw-tooth patterns in their host introns as RS sites.

**Fig. 2.**
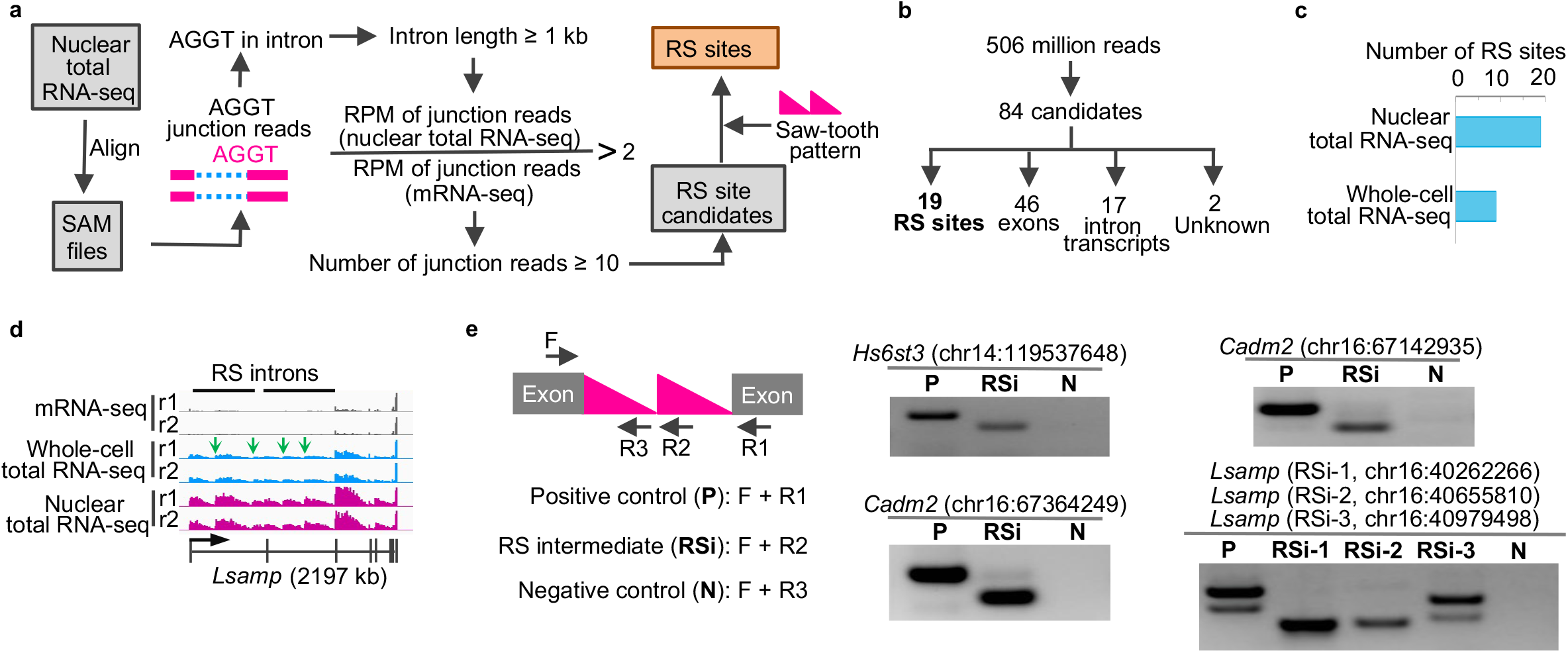
Identify RS sites using nuclear total RNA-seq data. (**a**) Schematic of the pipeline using nuclear total RNA-seq data to identify RS sites. (**b**) The 84 RS candidates. (**c**) Bar plot of RS sites identified using the two sequencing methods in the mouse cerebral cortex. (**d**) Sequencing profile at *Lsamp* locus. Green arrows indicate the four novel RS sites. **(e)** Using RT-PCR to detect RS splicing intermediates. Left, schematic of the primer design. Right, RT-PCR and gel results. Note, there are two bands in the positive control and RSi-3 in *Lsamp*, because there is an alternative splicing exon (60 bp) located between RS site 2 and RS site 3 in *Lsamp*.

### RS is a rare event in the mouse brain

By applying the pipeline to the 506 million uniquely mapped reads from our nuclear total RNA-seq data, we identified 84 RS site candidates (Supplementary Figure S1b). To further refine these candidates, we manually inspected the 84 sites, including the profiles of all sequencing reads and the junction reads at these genomic loci. We found that 19 of the 84 candidates are RS events that show a saw-tooth pattern (Fig. 2b,c and Supplementary Figure S1b). For the other candidates, 46 of them are exons (Supplementary Figure S1b,c) that were not annotated in the Ensembl release 93 database; 17 of them are likely nascent transcripts in introns (Supplementary Figure S1b, c); and 2 of them are unknown sites in the introns of *Etl4* and *Nufipl* (Supplementary Figure S1b).

The 19 RS sites include all the ten known RS sites in mice(5) (Supplementary Figure S1d). Nine RS sites (47%) we identified are novel in mice, including the four sites in the introns of *Lsamp* (Fig. 2d, green arrows). To validate the identified RS sites, we conducted RT-PCR to detect the RS splicing intermediates for three known RS sites in *Hs6st3* and *Cadm2* and three novel RS sites in *Lsamp* (Fig. 2e). Notably, we detected RS splicing intermediates for all the six RS sites (Fig. 2e), which independently supported the RS sites that we identified using the nuclear total RNA-seq data.

We next applied the same pipeline to the whole-cell total RNA-seq data and identified nine RS sites (Supplementary Figure S1d). We found that using nuclear total RNA-seq identified two-fold more RS sites than using whole-cell total RNA-seq (Fig. 2c), suggesting that nuclear total RNA-seq is an efficient approach to identify RS events.

To determine whether our criteria were too stringent, we lowered the cutoff of the counts of RS junction reads from 10 to five. We identified 103 additional RS site candidates (Supplementary Figure S1e). However, none of them exhibited saw-tooth patterns in the host introns, suggesting that lowering the cutoff of the counts of RS junction reads did not identify additional RS sites. Together, the finding that only 19 RS sites were identified from 506 million uniquely mapped reads indicates that RS is a rare event in the mouse brain.

### RS is restricted to the AGGT motif

Although GT is the most highly used motif for 5’ splice sites (5’SS), other motifs can also be used for 5’SS. To determine whether RS occurs in AGNN motifs where NN is not GT, we used a pipeline to identify RS sites in AGNN motif. This pipeline is similar to the one in Fig. 2a. Briefly, we extracted all junction reads spanning the AGNN sites, focused on the AGNN sites located in introns, selected AGNN sites that show more than two-fold enrichment of RS junction reads in nuclear total RNA-seq than in mRNA-seq, and then selected sites containing 10 or more RS junction reads as RS site candidates. As a result, we identified 405 AGNN RS site candidates (Supplementary Figure S2). However, visual inspection revealed that none of them exhibit saw-tooth patterns in the host introns. Thus, these results indicate that RS is restricted to the AGGT motif.

### RS is constitutive between cell types and sexes in the mouse cerebral cortex

RS has been shown to be constitutive in Drosophila and cell-type-specific in humans(3, 9). To determine the cell-type specificity of RS in mouse brain, we focused on two types of neurons, the excitatory neurons and the inhibitory neurons, which account for 85% and 15% of the neurons in the mouse cerebral cortex(12). We recently developed a genetic approach to tag and isolate cell-type-specific nuclei from the mouse brain and profiled gene expression in the excitatory and inhibitory neurons using nuclear total RNA-seq(12). We analyzed the nuclear total RNA-seq data from the two neuronal cell types (Supplementary Figure S1a). We applied our pipeline to these data and identified 17 RS sites in excitatory neurons and 18 RS sites in inhibitory neurons (Fig. 3a and Supplementary Figure S1d). Notably, all but one of the RS sites are common in both neuronal cell types (Fig. 3a). The only RS site unique to inhibitory neurons resides in the *Kcnip1* gene, which is only expressed in the inhibitory neurons (Fig. 3b and 3c). Thus, these results indicate that RS is largely constitutive between the two types of neurons in the mouse cerebral cortex.

**Fig. 3.**
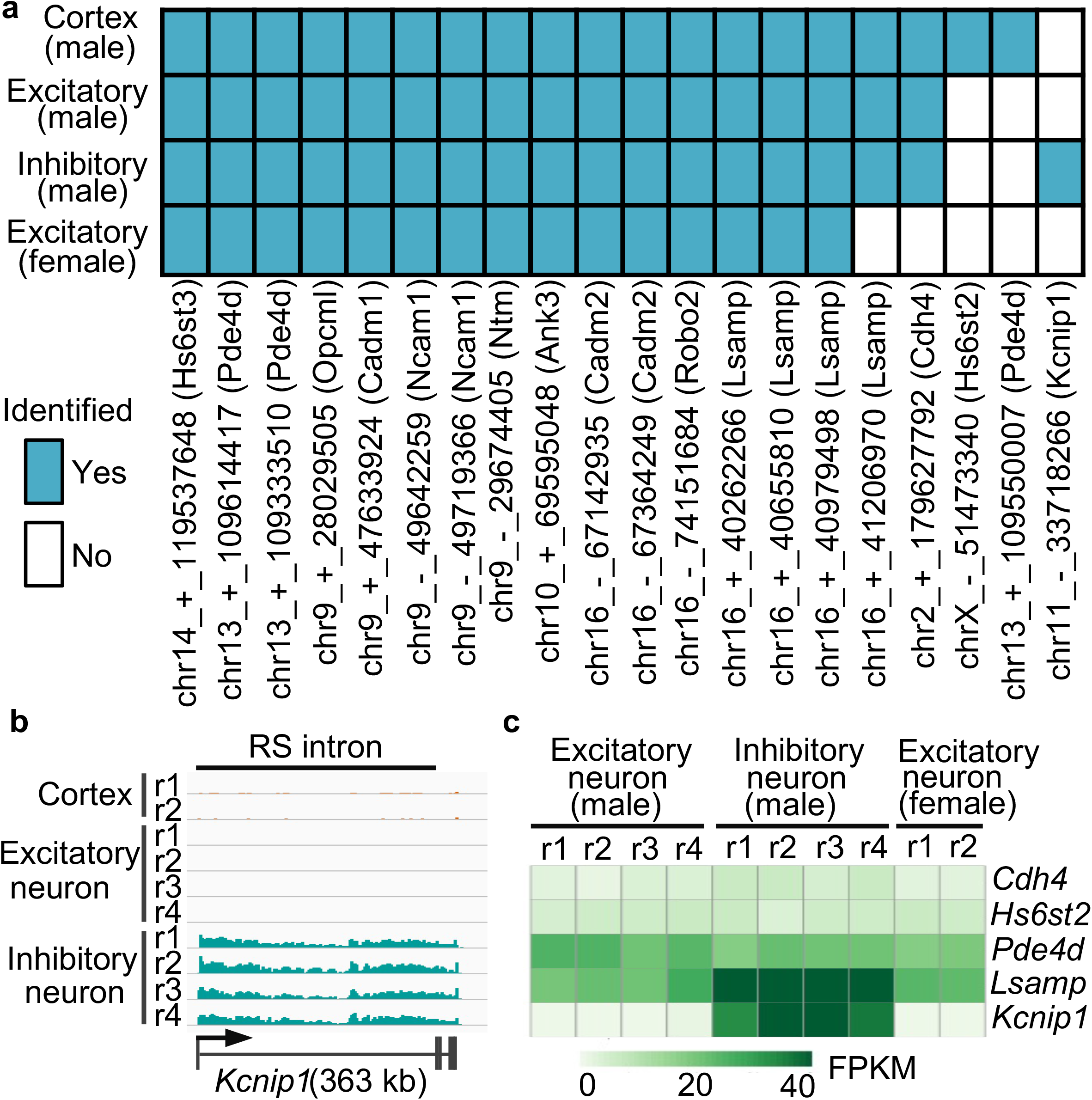
Cell-type and sex specificity of RS in the mouse cortex. **(a)** Heatmap of RS sites identified in male cortex, male cortical excitatory and inhibitory neurons, and female cortical excitatory neurons. **(b)** Nuclear total RNA-seq profile (male) at *Kcnip1* locus. **(c)** Heatmap of expression levels of five genes in three cell types. FPKM, fragment per million uniquely mapped reads per kilobase of exonic region.

We recently also profiled gene expression in the excitatory neurons in the cerebral cortex of female mice using nuclear total RNA-seq(12). To investigate the sex specificity of RS, we analyzed the nuclear total RNA-seq data of excitatory neurons from the cerebral cortex of female mice(12). We identified 15 RS sites (Supplementary Figure S1d), which are all included in the 17 RS sites we identified from male excitatory neurons (Fig. 3a). The remaining two RS sites are unlikely male specific, because we also identified seven and five junction reads for them in the female data (Supplementary Figure S1d), although they failed to pass our criteria of 10 junction reads. Together, these results indicate that RS is constitutive between male and female excitatory neurons in the mouse cerebral cortex.

### Characteristics of RS sites

In consistent with previous reports, RS sites are highly conserved (Supplementary Figure S3a), are located in long introns (Supplementary Figure S3b), prefer the first two introns (Supplementary Figure S3c, 75% in first intron and 25% in second intron), and are located in genes that are specifically expressed in the brain (Supplementary Figure S3d). Furthermore, compared to non-RS AGGT sites, RS AGGT sites are enriched for AGGTAAGT motif that complements with the 5’ conserved sequence of U1 snRNA (Supplementary Figure S3e-g), are enriched for thymine and cytosine in the 20 nt regions upstream of RS AGGT sites (polypyrimidine tract, Supplementary Figure S3h), and show higher 3’ splice site scores when examined using MaxEntScan(16) (Supplementary Figure S3i).

### Splicing intermediates linking RS exons to the downstream annotated exons

RS uses exon definition mechanism(5, 7), thus suggesting a possibility of splicing intermediates linking RS exons to the downstream annotated exons (downstream splicing intermediates, Fig. 4a). Although Sibley et al. reported that RS exons were not detectable in mRNA transcripts and were included only after blocking the 5’SS using antisense oligonucleotide(5), the extent to which the downstream splicing intermediates exist in physiological conditions remains largely unexplored.

**Fig. 4.**
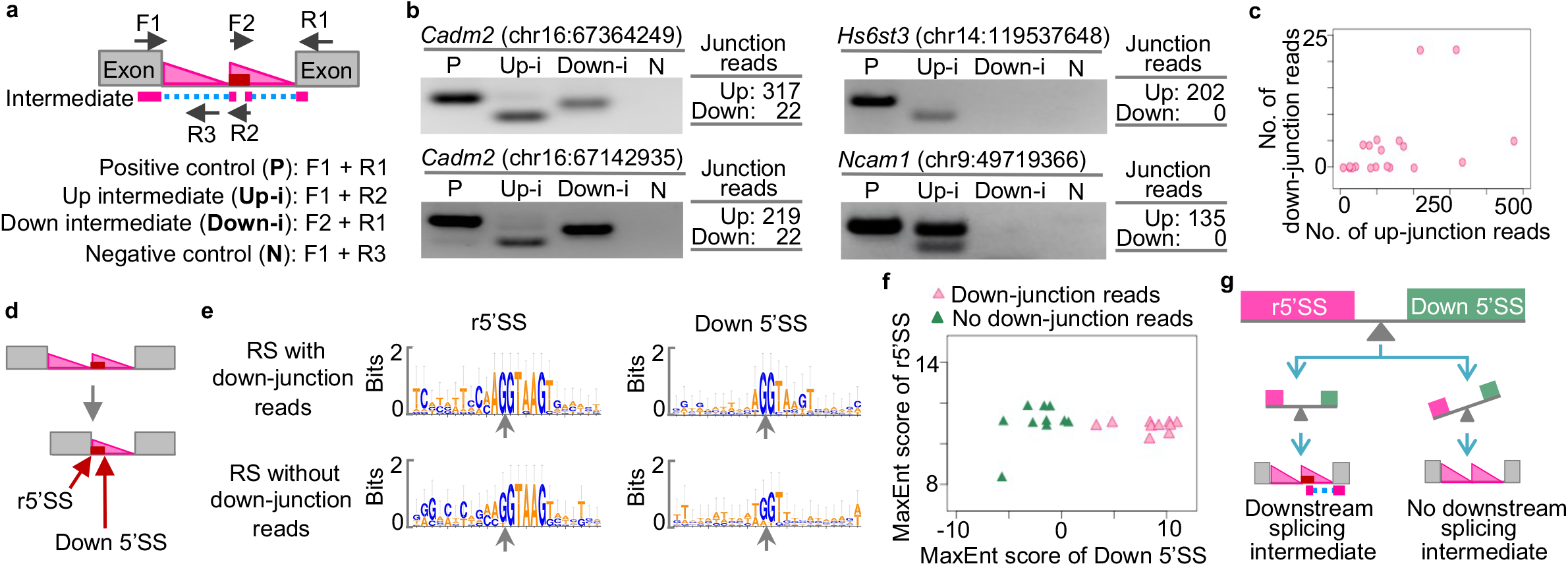
Downstream RS splicing intermediates. **(a)** Schematic of the up- and down-junction reads that represent the upstream and downstream RS splicing intermediates and the primer design. **(b)** Validation of the upstream (Up-i) and downstream (Down-i) RS splicing intermediates using RT-PCR. P, positive control. N, negative control. Left, RT-PCR and gel results. Right, numbers of junction reads from the nuclear total RNA-seq data. **(c)** Scatter plot of the numbers of up- and down-junction reads of the 20 RS sites. **(d)** The reconstituted 5’SS (r5’SS) and the downstream 5’SS (Down 5’SS). **(e)** Sequence logos of the r5’SS and the Down 5’SS of RS sites with or without down-junction reads. **(f)** Scatter plot of the MaxEnt scores of r5’SS and Down 5’SS of RS sites. **(g)** Model of the 5’SS strengths and the downstream splicing intermediates.

To identify downstream splicing intermediates of RS sites, we developed a computational pipeline to identify junction reads that link RS exons to downstream annotated exons (down-junction reads, Fig. 4a). Briefly, we obtained all the junction reads from our nuclear total RNA-seq data and then scanned the 350 nt region downstream of the RS AGGT site for junction reads that link to the downstream exons. We found downjunction reads for 10 RS sites, but not for the other 10 sites (Supplementary Figure S4a). To validate the findings of down-junction reads, we conducted RT-PCR to detect the downstream splicing intermediates for four RS sites (Fig. 4b). Notably, for the two RS sites that show both up- and down-junction reads, we detected both upstream and downstream splicing intermediates (Fig. 4b, left panel). However, for the two RS sites that only show up-junction reads, we only detected the upstream splicing intermediates, not the downstream splicing intermediates (Fig. 4b, right panel). Thus, the RT-PCR results are consistent with the results from the nuclear total RNA-seq. Together, these results indicate that ten RS sites have downstream splicing intermediates, while the other ten sites do not have downstream splicing intermediates.

### Downstream splicing intermediates are associated with high strengths of the 5’ splice sites at the 3’ end of the RS exons

To understand why some RS sites do not show downstream splicing intermediates, we first asked whether these RS sites also show low abundance of upstream splicing intermediates. We analyzed the numbers of up- and down-junction reads for the 20 RS sites. However, we found no positive correlation between the two (Fig. 4c), suggesting that the absence of downstream splicing intermediates for the 10 RS sites is not associated with the low abundance of upstream splicing intermediates.

Previous studies have showed that the competition between the two 5’SS was associated with the inclusion of the RS exons in mature transcripts(5, 7, 9). Thus, another hypothesis is that the 5’SS at the two ends of the RS exons, the reconstituted 5’SS (r5’SS) after the first step of splicing and the downstream 5’SS (Down 5’SS) (Fig. 4d), affect the existence of the downstream splicing intermediates. To test this hypothesis, we quantified the strengths of r5’SS and Down 5’SS using MaxEntScan(16). We found that RS sites with down-junction reads show higher MaxEnt scores of Down 5’SS compared to RS sites without down-junction reads (*P* = 4.28^-9^, one-tailed t-test) (Fig. 4e,f and Supplementary Figure S4a). By contrast, the scores of r5’SS are comparable between the two groups (Fig. 4e,f and Supplementary Figure S4a). Together, these results support a model that the downstream splicing intermediates of RS sites are likely associated with the high strengths of 5’SS at the 3’ end of the RS exons (Fig. 4g).

### Cryptic exons use a RS-like splicing mechanism

During our analysis of RS site candidates, we found genomic loci that show exon-like splicing patterns (Fig. 2b). Some of these loci were later annotated as exons in Ensembl database, while the others are not. We termed these unannotated genomic loci as RS-like cryptic exons (Fig. 5a). To systematically identify RS-like cryptic exons, we developed a computational pipeline (Supplementary Figure S4b). To exclude unannotated exons from our downstream analysis, we used a criterion that the normalized count of up-junction reads in nuclear total RNA-seq should be higher than that in mRNA-seq. By applying this pipeline to our nuclear total RNA-seq data, we identified 22 RS-like cryptic exons in the introns of 21 long genes (Supplementary Figure S4c). To validate the RS-like cryptic exons, we conducted RT-PCR to detect splicing intermediates for three RS-like cryptic exons. We confirmed the upstream and downstream splicing intermediates for all the three RS-like cryptic exons (Fig. 5b). Notably, we found that although the cryptic exons in *Lrrc4c* and *Magi1* were not included in the mature mRNA, the cryptic exon in *Nova1* was included in a part of the mature mRNA (Fig. 5b), suggesting that RS-like cryptic exons may be included in the mature mRNA.

**Fig. 5.**
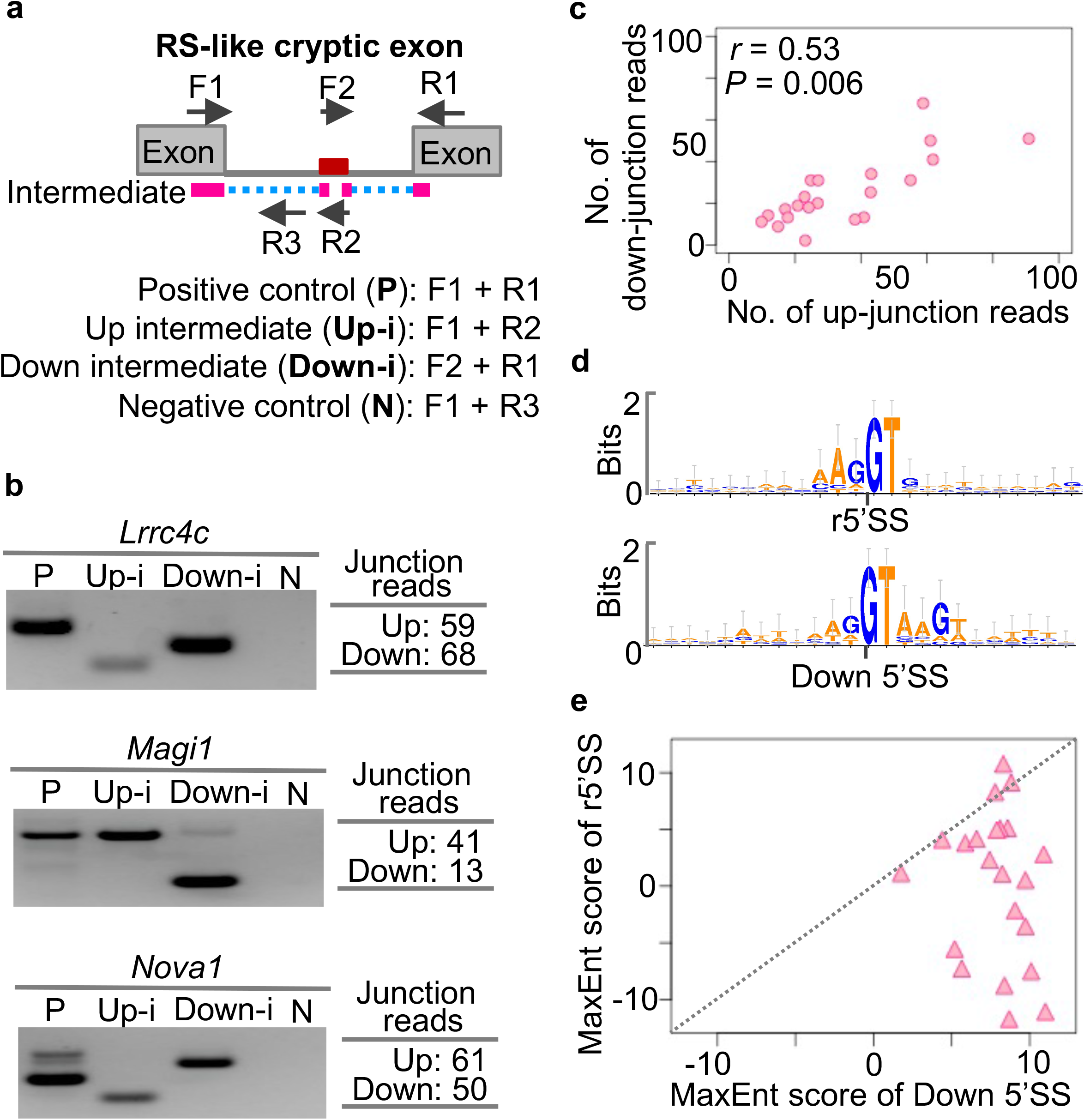
Cryptic exons use a RS-like splicing mechanism. **(a)** Schematic of RS-like cryptic exons and the primer design. **(b)** Validation of the upstream (Up-i) and downstream (Down-i) splicing intermediates using RT-PCR. P, positive control. N, negative control. Left, RT-PCR and gel results. Right, numbers of junction reads from the nuclear total RNA-seq data. **(c)** Scatter plot of the numbers of up- and down-junction reads of the RS-like cryptic exons. **(d)** Sequence logos of the r5’SS and Down 5’SS of the RS-like cryptic exons. **(e)** Scatter plot of the MaxEnt scores of the r5’SS and Down 5’SS of the RS-like cryptic exons.

To identify features of RS-like cryptic exons, we analyzed the correlation between the up- and down-junction reads for them. We found that RS-like cryptic exons exhibit a positive correlation between the numbers of up- and down-junction reads (Pearson correlation coefficient *r* = 0.53, *P-value* = 0.01, Fig. 5c), which contrasts with the results of RS sites (Fig. 4c). To investigate the strengths of the 5’SS at the two ends of RS-like cryptic exons, we examined the sequence compositions of the r5’SS and Down 5’SS using WebLogo(17). We found that the RS-like cryptic exons show an enrichment of 5’SS motif (AGGTAAGT) at the Down 5’SS but not at the r5’SS (Fig. 5d). Furthermore, we quantified the strengths of r5’SS and Down 5’SS using MaxEntScan(16) and found that the MaxEnt scores of r5’SS are significantly lower than that of Down 5’SS in RS-like cryptic exons (*P-value* < 0.0001, one-tailed t-test, Fig. 5e). One exception is the RS-like cryptic exon in *Magi1*, which shows a much higher MaxEnt score at the r5’SS than the Down 5’SS (Fig. 5e and Supplementary Figure S4c). Notably, the *Magi1* intron exhibits a weak saw-tooth pattern (Supplementary Figure S4d), which failed to pass the stringent criteria in our identification of RS sites. Taken together, these results demonstrate that some cryptic exons may use an RS-like mechanism for splicing.

## Discussion

In this study, we demonstrated that RS is a rare event in the mouse brain and that nuclear total RNA-seq is an efficient approach to investigate RS events. We developed a novel pipeline to identify RS sites, determined the cell-type and sex specificity of RS, and characterized the genomic features of RS sites. Through analysis of the primary sequences, we found that RS splicing intermediates are associated with the primary sequence context. We also discovered that some cryptic exons use an RS-like mechanism for splicing.

The extent to which RS is used for intron splicing in mammals remains inconclusive. Here we analyzed high-depth nuclear total RNA-seq data and identified 20 RS sites in mice, which is consistent with previous studies(3, 5) that RS is a rare splicing process for a small group of long introns in the mammalian genome. We found that whether or not including the saw-tooth patterns to define RS sites might be the major reason for the differences of the reported numbers of RS events(3, 5, 9, 11). For example, we identified 65 AGGT sites and 405 AGNN sites that have 10 or more junction reads, but they did not show saw-tooth patterns in the host introns. Therefore, we did not consider them as RS sites. Notably, the relationship between saw-tooth patterns and RS events remains unclear. Further studies are needed to investigate whether the saw-tooth pattern should be a required criterion to define an RS event.

Long introns exhibit a high rate of creating new exons during evolution(18), but the underlying mechanisms are not fully understood. Our discovery of RS-like cryptic exons indicates that long introns may acquire novel exons via the RS-like cryptic exons. This is supported by the findings that more than 6000 human annotated exons are RS-like exons(19). In addition, the numbers of RS sites in *Drosophila melanogaster* are about 15 times more than that in humans(3, 5, 7–9), but the numbers of RS-like annotated exons in *Drosophila* are 2~100 times less than that in humans(5, 7, 19). Thus, future studies are needed to investigate RS, RS-like cryptic exons, and RS-like annotated exons in evolutionarily distinct species to determine their associations during evolution.

RS genes are genetically linked to various human brain disorders. For example, *ANK3* encodes ankyrin-G and is linked to autism spectrum disorders, attention deficit hyperactivity disorder, intellectual disability, and bipolar disorder(20–23). Also, *NTM* encodes neurotrimin and is linked to autism spectrum disorders and attention deficit hyperactivity disorder(24, 25). The *PDE4D* encodes phosphodiesterase 4D and is linked to schizophrenia, psychosis, acrodysostosis, and neuroticism(26–28). Notably, PDE4D Inhibitors are in clinical trials for the treatment of Alzheimer’s disease and Fragile X syndrome(29, 30). Given that the disruption of the RS process interfered with the RS gene function and caused abnormality in the central nervous system(7), further investigation will be necessary to illuminate whether RS mechanism contributes to the pathophysiology of these human brain disorders.

## Materials and Methods

### Animals

The C57BL/6 mice were obtained from the Jackson Laboratory. Mice were maintained in the same genetic background and were kept on a regular 12-hour light/12-hour dark cycle. Animal studies were carried out in accordance with the National Institutes of Health’s Guide for the Care and Use of Laboratory Animals recommendations. The study protocol was approved by the Institutional Animal Care and Use Committee (IACUC) of the New York Institute of Technology. The study is reported in accordance with the ARRIVE guidelines (https://arriveguidelines.org).

### RNA isolation and RT-PCR

The brain was dissected from 8-week-old mice. Total RNA was extracted from the brain tissues using TRIzol reagent (Invitrogen, #15596026) according to the manufacturer’s instructions. The integrity and quality of the isolated RNA were determined by the Nanodrop and the Bioanalyzer. The Prime script reverse transcript kit (Takara, #RR037A) was used to synthesize the cDNA. The RT-PCR experiments were conducted using the GoTaq G2 Hot Start Master mix (Promega #M7422). The PCR primers were listed in the Supplementary Figure S5. The RT-PCR were conducted using the following setting: 94 °C for 3 minutes, followed by 32-40 cycles of thermocycling (94 °C for 30 seconds, 52 °C-56 °C for 30 seconds, 72 °C for 30 seconds), and 72 °C for 3 minutes.

### Statistical analysis

All statistical analyses were performed in the R software version 3.6.1 (https://www.r-project.org/).

### Nuclear total RNA-seq data analysis

The analysis pipeline was described previously(12, 13, 31, 32). In brief, raw data in sra files were downloaded from the EBI European Nucleotide Archive database(33) using the accession numbers listed in Figure S1. The fastq-dump.2.9.6 of NCBI SRA ToolKit was used to extract the FASTQ files using the parameter of “--split-3”. STAR(34) was used to map the FASTQ raw reads into mouse mm10 genome using the parameters of “--runThreadN 40 --outFilterMultimapNmax 1 -- outFilterMismatchNmax 3”. The samtools view(35) was used to convert sam files into bam files. The samtools sort was used to sort the bam files. The samtools index was used to index the sorted bam files. The bamCoverage(36) was used to convert the sorted bam files into strand-specific bigwig files. The bamCoverage parameters that were used included “--filterRNAstrand forward --binSize 1 -p 40 -o” for plus strand and “-- filterRNAstrand reverse --binSize 1 -p 40 -o” for minus strand.

### Sequencing data visualization

All sequencing data, including the bigwig files and bam files, were visualized in the IGV_2.8.2 genome browser(37).

### Genome annotation

The gtf file of mouse genome annotation was downloaded from the Ensembl release 93(38).

### Junction reads

A read pair is considered as a junction read if its CIGAR in sam files contains “N”. Junction reads were extracted from sam files and were saved into a junctionread-specific sam files. These sam files were further converted into bam files using samtools. The junction-read-specific bam files were loaded into IGV for visualization.

### Junction reads spanning AGGT sites

All AGGT sites (20,403,114) in the mouse mm10 genome were identified, and only AGGT sites (4,767,575) located in the gene sense regions were kept for further analysis. The AGGT sites that were kept were used to screen the junction-read-specific sam files. The numbers of junction reads spanning each AGGT site (joining the upstream exon and sequences following GT) were counted. The counts of junction reads were further normalized to the sequencing depth to obtain the RPM values.

### Pipeline to identify RS sites

The schematic of this pipeline is shown in Fig. 2a. Briefly, AGGT sites located in introns longer than or equal to 1 kb were extracted. Sites that showed a larger than two-fold RPM value of junction reads in total RNA-seq data than in mRNA-seq data were kept for downstream analyses. The counts of junction reads of biological replicates were merged, and AGGT sites that contained 10 or more junction reads were identified as RS site candidates. The RS site candidates were further refined as RS sites if the host intron showed a clear saw-tooth pattern by visual inspection.

### FPKM of nuclear total RNA-seq data

The number of reads mapped to the exonic regions of each gene were calculated to get the raw counts. The raw counts were then normalized to the exon lengths of that gene and to the sequencing depth of that data set to get the FPKM values.

### Phylogenetic p-value (phyloP) scores

The phyloP scores, which were calculated by the PHAST package for multiple alignments of 59 vertebrate genomes to the mouse genome, were obtained from the UCSC Genome Browser (http://hgdownload.cse.ucsc.edu/goldenpath/mm10/phyloP60way/).

### Gene expression profiles in 22 mouse tissues

The expression profiles (FPKM values) of RS genes in 22 mouse tissues were obtained from the LongGeneDB database (https://longgenedb.com).

### WebLogo analysis

WebLogo 3(17) (http://weblogo.threeplusone.com) was used to perform the sequence logo analysis. The Output Format was chosen as “PNG (high res.)”, and the Stacks per Line was set to “80”. The default values were used for other parameters.

### MaxEntScan 3’SS analysis

The 3’SS scores were calculated by MaxEntScan::score3ss (http://hollywood.mit.edu/burgelab/maxent/Xmaxentscan_scoreseq_acc.html)(16). The input sequences were composed of the 18 nt region upstream of the AGGT, the AGGT motif, and the one nucleotide following AGGT (18nt + AGGT + 1nt). The three models - Maximum Entropy Model, First-order Markov Model, and Weight Matrix Model - were selected. The MaxEnt scores were used as the 3’SS scores.

### Reconstituted 5’ splice sites (r5’SS)

The r5’SS sequences were composed of the last 30 nucleotides of the upstream exon, the GT motif, and the 20 nucleotides following AGGT (30nt + GT + 20nt).

### MaxEntScan 5’SS analysis

The 5’SS scores were calculated by MaxEntScan::score5ss (http://hollywood.mit.edu/burgelab/maxent/Xmaxentscan_scoreseq.html)(16). The input sequences for 5’SS were composed of three nucleotides before the GT, the GT motif, and the four nucleotides following AGGT (3nt + GT + 4nt). The input sequences of r5’SS and Down 5’SS were listed in Supplementary Figure S4a and S4c.

### Pipeline to identify RS-like cryptic exons

The schematic of this pipeline is shown in Supplementary Figure S4b. Briefly, AGGT sites located in introns longer than or equal to 50 kb were extracted. The AGGT sites that showed a larger RPM value of junction reads in total RNA-seq data than in mRNA-seq data were kept for downstream analyses. The counts of the junction reads of the biological replicates were merged. AGGT sites contained 10 or more up-junction reads and two or more down-junction reads were identified as candidates of RS-like cryptic exons. The RS-like cryptic exon candidates were further refined as RS-like cryptic exons if the host intron showed an exon-like but not saw-tooth like pattern.

## Data availability

The data sets supporting the conclusions of this article are available in the NCBI GEO database with the accession numbers listed in Supplementary Figure S1a. The custom code supporting the conclusions of this article is available in the GitHub repository, https://github.com/Jerry-Zhao/RS2020.

## Author contributions

SM and YTZ conceived the project, designed the experiments, curated data, and wrote the manuscript. SM performed the experiments. YTZ performed the computational analyses. All authors read and approved the final manuscript.

## Acknowledgements

We thank Dr. Raddy Ramos, Dr. Weikang Cai, and members of the Zhao Laboratory for helpful discussions and comments on the manuscript. We thank the Center for Biomedical Innovation at the New York Institute of Technology College of Osteopathic Medicine for support.

## Supporting information

**Supplementary Figure S1:**
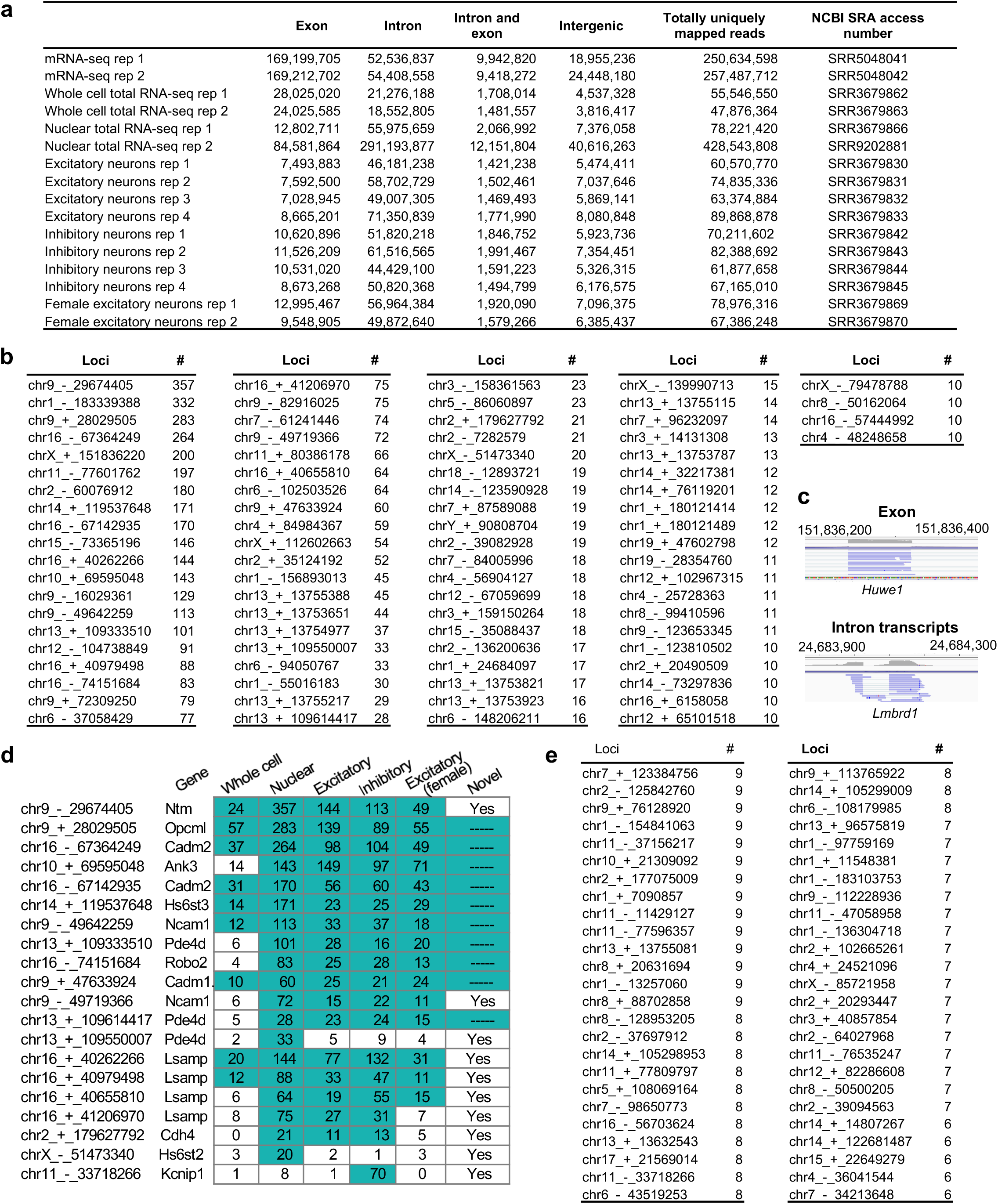
Novel RS sites. (**a**) The mapping statistics and access numbers of RNA-seq data utilized in this study. (**b**) The loci and numbers of junction reads (#) of the 84 RS site candidates. (**c**) Snapshots of the junction reads at the Huwe1 and Lmbrd1 RS candidate sites. (**d**) Heatmap of numbers of RS junction reads at each RS site in different total RNA-seq data sets. A green box indicates that the RS site was identified in that data set. Note, the enrichment of junction reads in whole cell data compared to that in mRNA-seq data is less than 2-fold for Ank3. (**e**) The loci and read numbers (#) of the top 50 additional RS site candidates when lowering the cutoff of junction read count from 10 to 5.

**Supplementary Figure S2:**
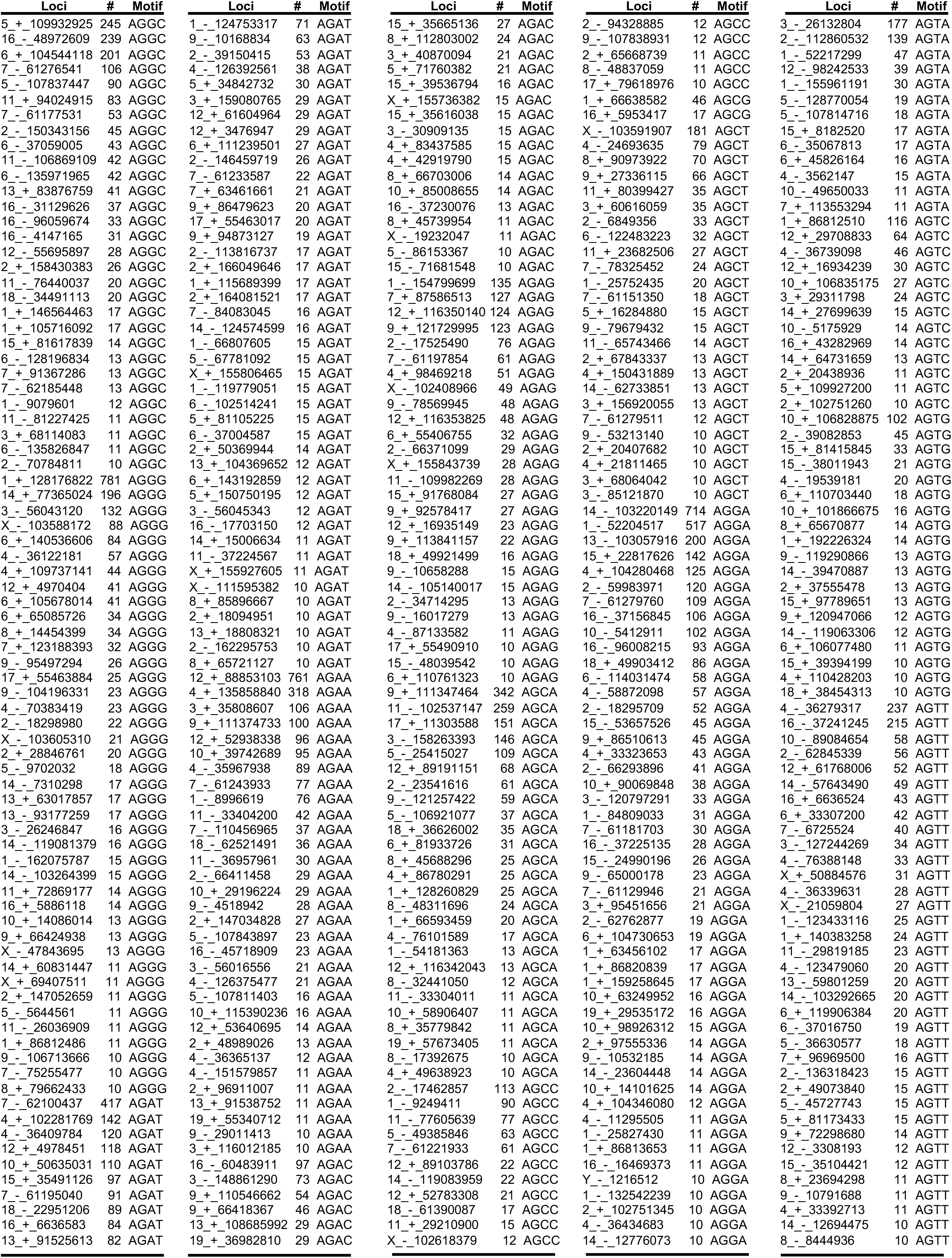
The 405 AGNN RS site candidates. #, number of junction reads.

**Supplementary Figure S3.**
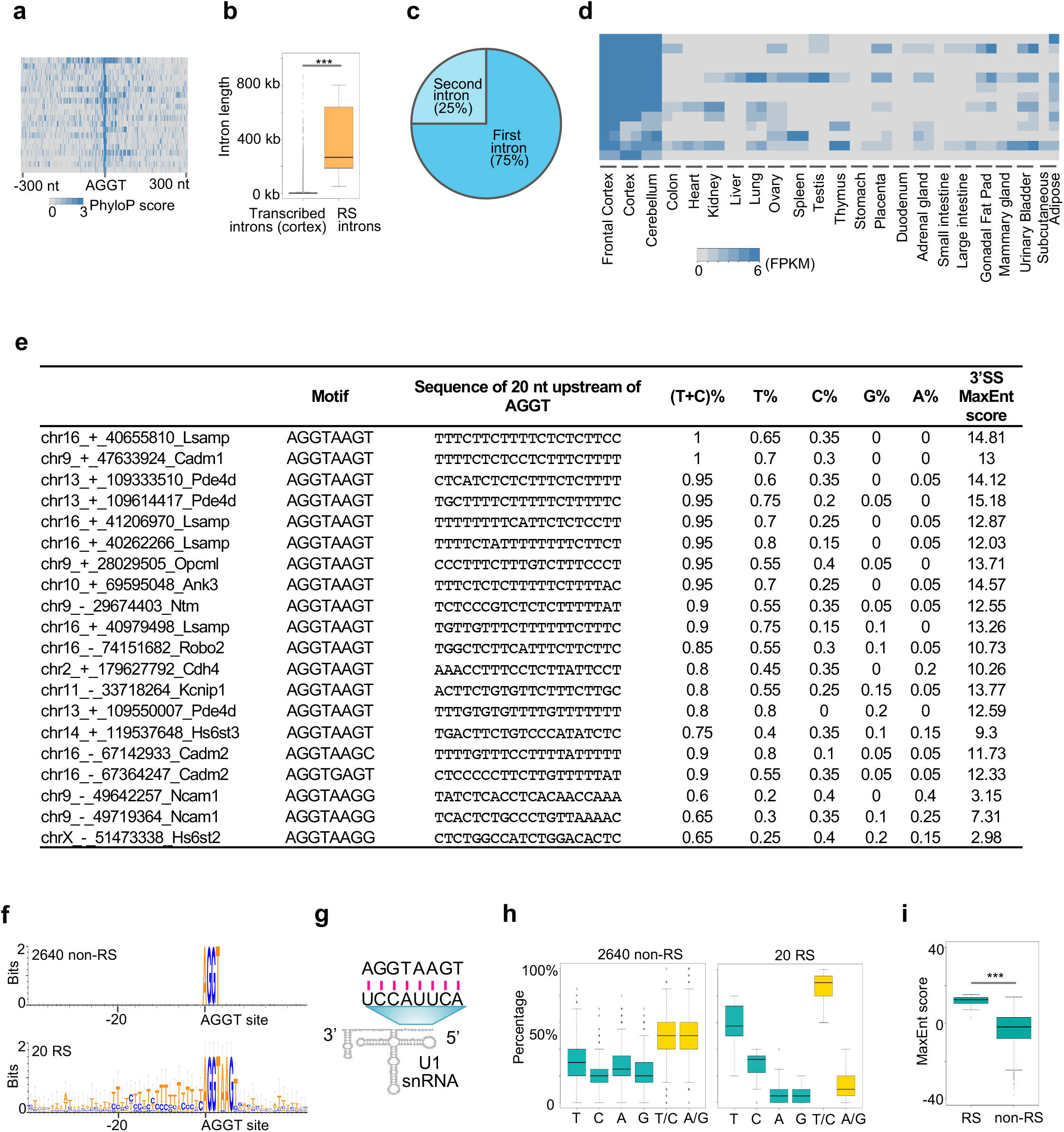
Characteristics of RS sites. (**a**) Heatmap of phyloP score of RS sites and the flanking regions. (**b**) Boxplot of lengths of RS introns and introns transcribed in the mouse cortex. ***, *P* < 0.0001, one-tailed t-test. (**c**) Pie chart of locations of RS introns in host genes. (**d**) Heatmap of expression levels of RS genes in 22 mouse tissues. (**e**) The sequence motifs, nucleotide percentages, and 3’SS MaxEnt scores of the 20 RS sites. (**f**) Sequence logos of the 64 nt regions surrounding the 2640 non-RS AGGT sites and the 20 RS AGGT sites. (**g**) Schematic of the sequence base pairing between the AGGTAAGT motif and U1 snRNA. (**h**) Boxplots of the percentages of nucleotides in the 20 nt region upstream of the 2640 non-RS AGGT sites and the 20 RS AGGT sites. (**i**) Boxplot of MaxEnt 3’ splice site (3’SS) scores of the 20 RS AGGT sites and the 2640 non-RS AGGT sites. ***, *P* < 0.0001, one-tailed t-test.

**Supplementary Figure S4.**
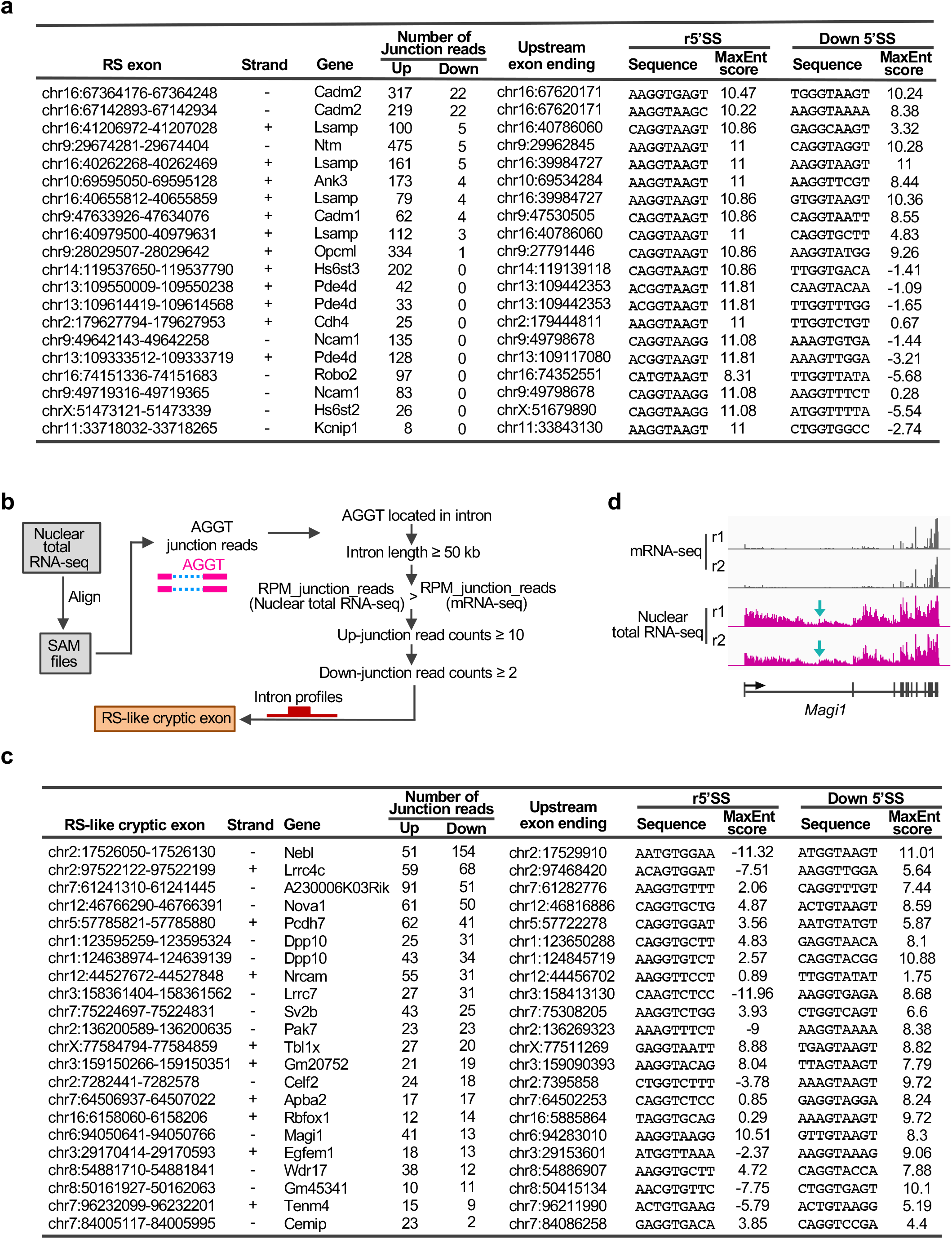
RS exons and RS-like cryptic exons. (**a**) The genomic loci, numbers of junction reads, 5’SS sequences, and 5’SS MaxEnt scores of RS exons. (**b**) Schematic of the pipeline utilizing nuclear total RNA-seq data to identify RS-like cryptic exons. (**c**) The genomic loci, numbers of junction reads, 5’SS sequences, and 5’SS MaxEnt scores of RS-like cryptic exons. (**d**) Sequencing profile at *Magi1* locus. Green arrows indicate the putative RS AGGT loci.

**Supplementary Figure S5.**
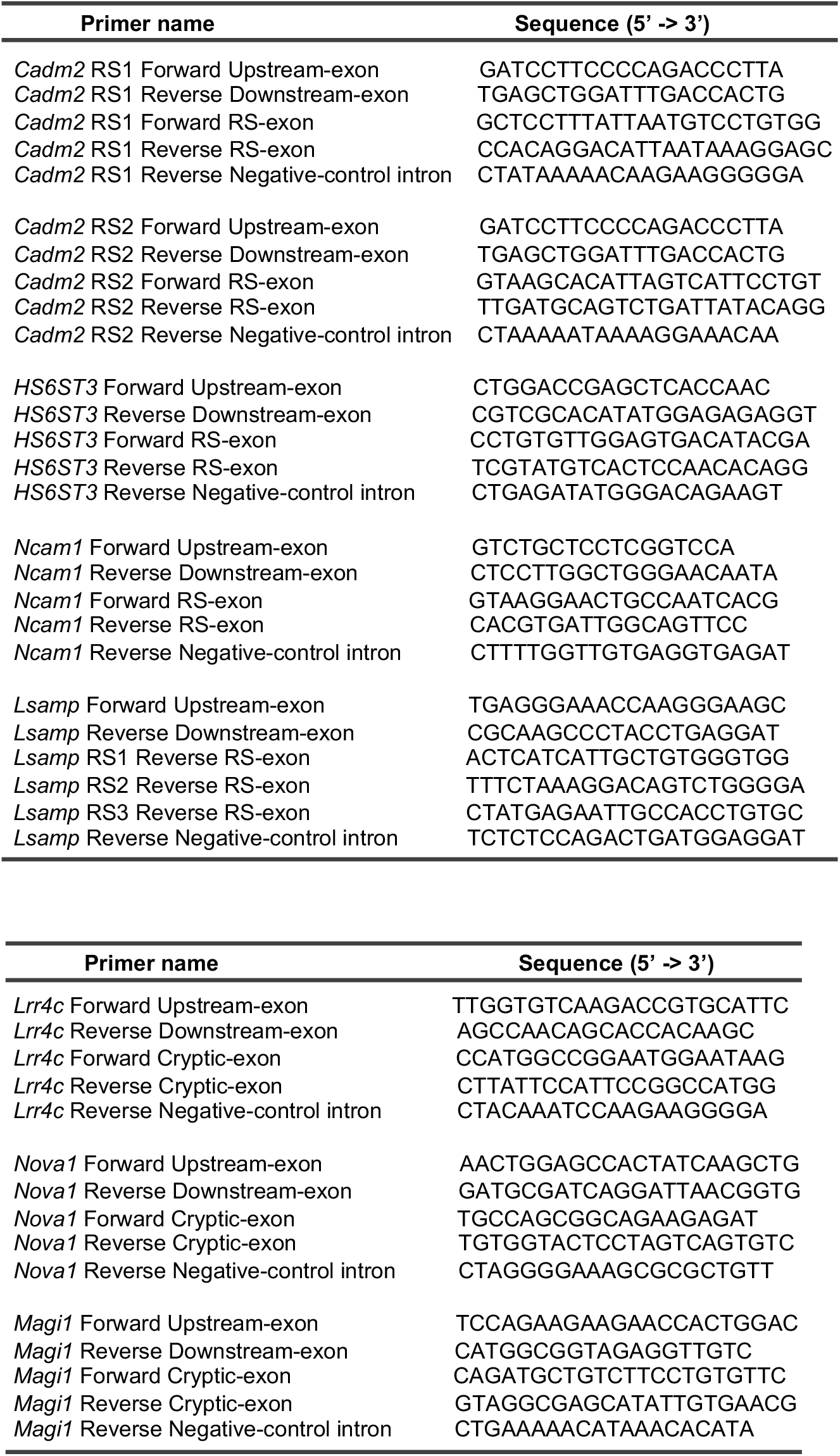
RT-PCR primer sequences for the RS sites (top table) and RS-like cryptic exons (bottom table). *Cadm2* RS1: chr16:67364249. *Cadm2* RS2: chr16:67142935. *Lsamp* RS1: chr16:40262266. *Lsamp* RS2: chr16:40655810. *Lsamp* RS3: chr16:40979498.

**Supplementary Figure S6.**
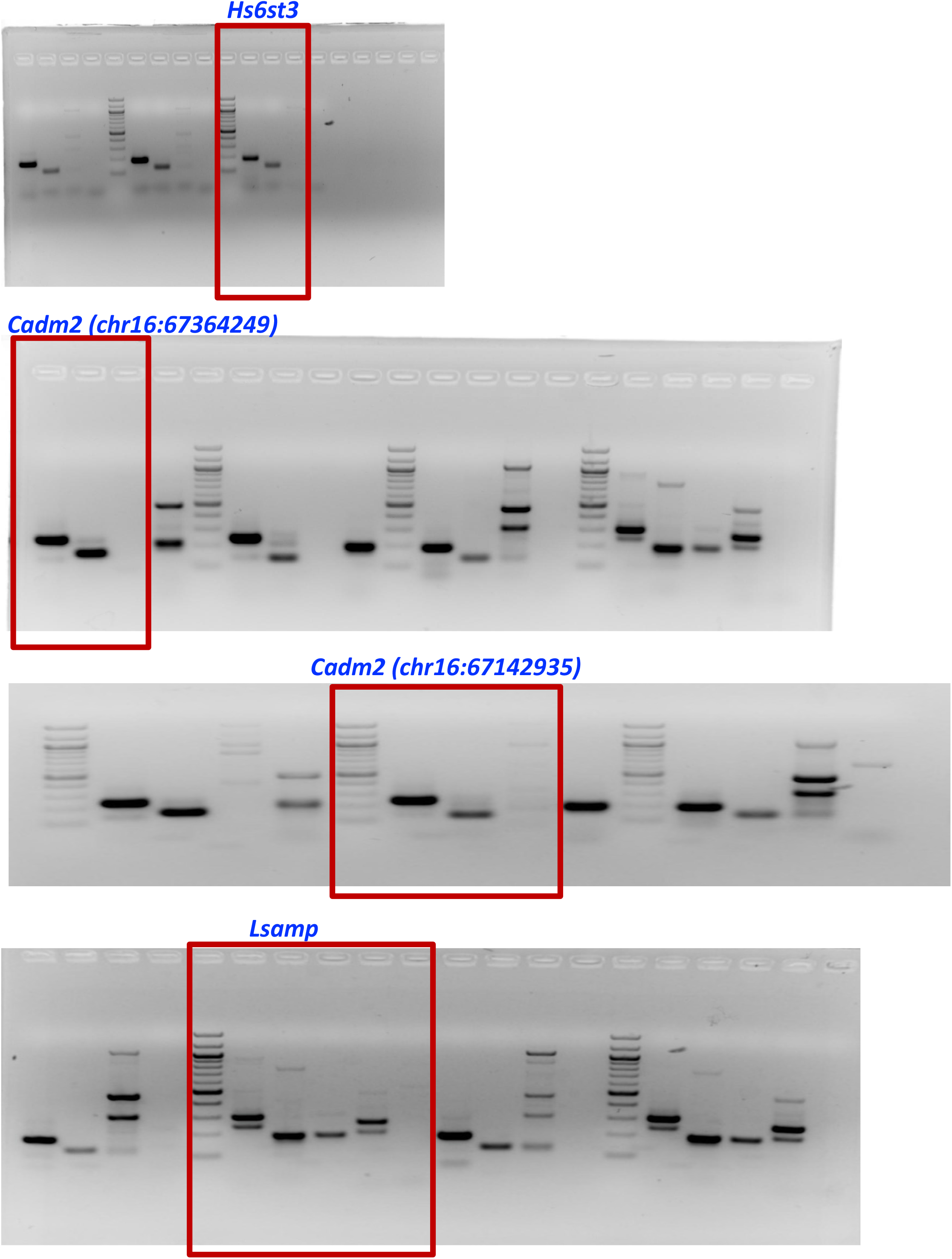
The full-length gels for plots in Figure 2.

**Supplementary Figure S7.**
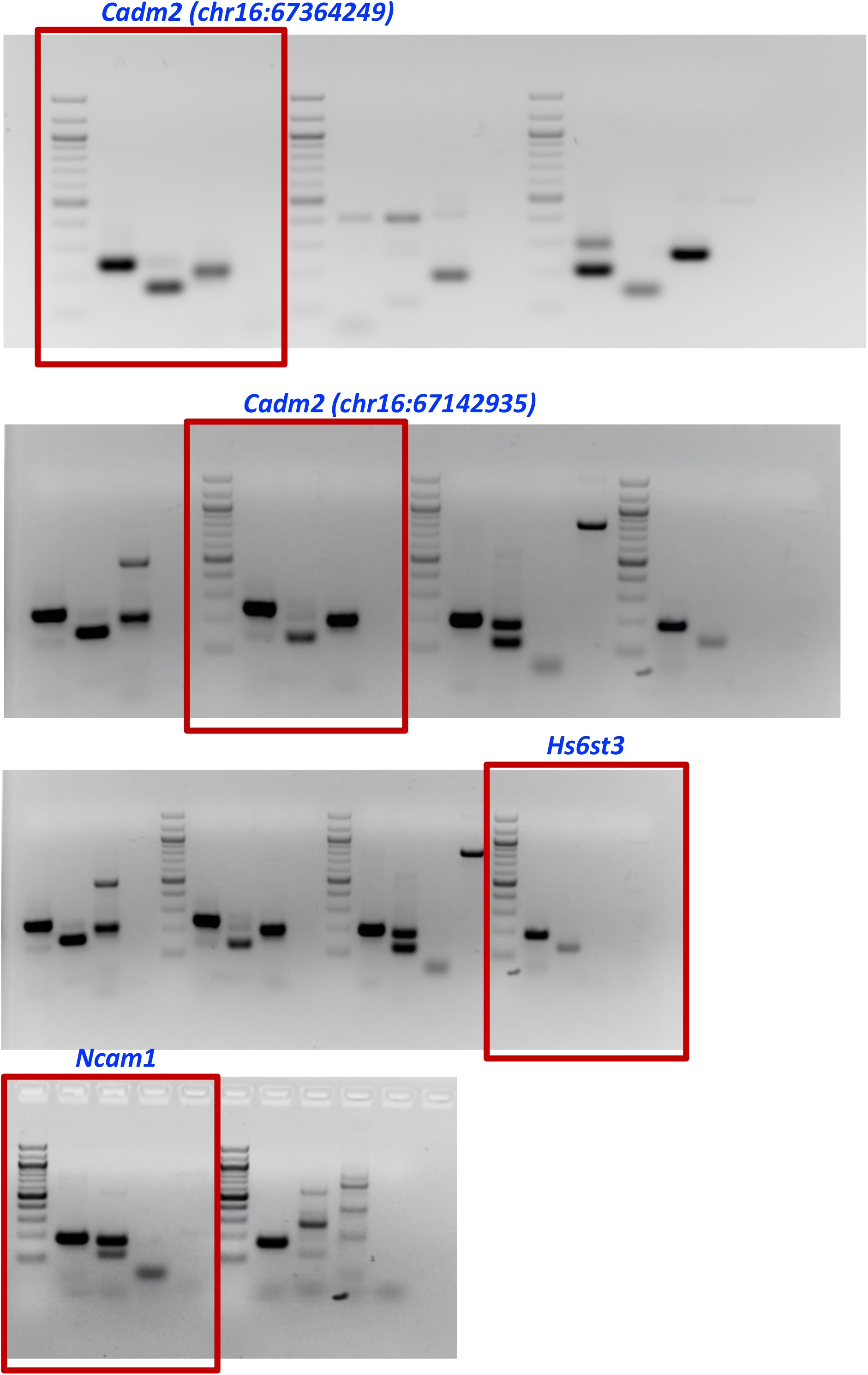
The full-length gels for plots in Figure 4.

**Supplementary Figure S8.**
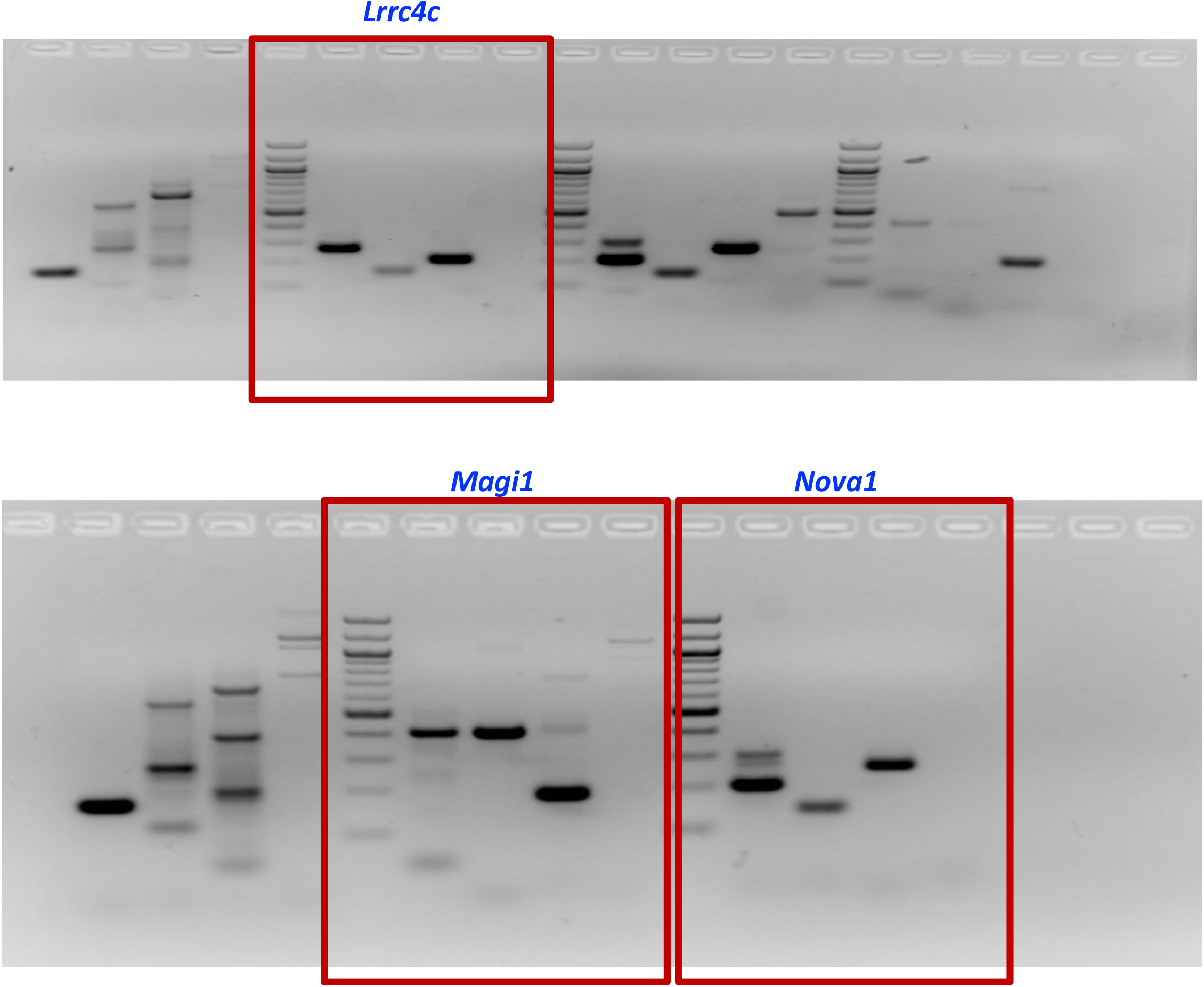
The full-length gels for plots in Figure 5.

